# Sequential acquisition of 1p31.1-p12 LOH and 1q Gain is a common double-hit event in relapsed/refractory myeloma

**DOI:** 10.64898/2026.05.19.726252

**Authors:** Naser Ansari-Pour, Sarah Gooding, Mohammad H Kazeroun, Seyed Alireza Hasheminasab, Evie Fitzsimons, Selina Chavda, Alessandro Laganà, Erin Flynt, Udo Oppermann, Karthik Ramasamy, Kwee Yong, Angela Hamblin, Jill Corre, Herve Avet-Loiseau, Nikhil Munshi, Mehmet Samur, Anjan Thakurta

## Abstract

Therapy-driven genomic changes in multiple myeloma (MM) remain poorly defined. We analyzed whole-genome sequencing (WGS) data from relapsed/refractory MM (rrMM, N=386) and identified regional 1p31.1–p12 (hereafter 1pCEN, a region proximal to the centromere) loss-of-heterozygosity (LOH) as the only enriched aberration showing strong therapy-associated clonal selection (clonal timing rank fold-change = 3.7, P<2.2×10^-16^). This event showed enriched co-occurrence with 1qGain (OR = 2.3 (1.5-3.8), P=2×10^-4^) forming a recurrent “double-hit” in rrMM. To validate the clonal selection process, we examined three longitudinal cohorts (180 patients, 390 samples) and confirmed clonal expansion of 1pCEN and consistent prevalence of the 1pCEN+1q double-hit (20–24%). Survival analyses demonstrated significantly reduced progression-free survival in rrMM patients with this double-hit compared with those without. Comparison with a large newly diagnosed MM (ndMM) cohort confirmed previously-described 1p32 LOH is the prognostic locus at baseline, whereas 1pCEN is therapy-selected and largely independent of the 1p32 locus. Thus, 1pCEN+1q represents a recurrent double-hit event that clonally emerges in rrMM, conferring selective advantage under drug exposure and is distinct from the ndMM high-risk markers defined by current consensus guidelines. These findings nominate 1pCEN as a new genomic biomarker in rrMM and 1pCEN+1q may help patient stratification for therapeutic monitoring.

**Key Points:** A therapy-driven common genomic double-hit (1p31.1–p12 LOH with 1q gain) clonally emerges in relapsed/refractory myeloma.

## Introduction

Multiple myeloma (MM) is a malignancy of clonal plasma cells characterized by marked genetic heterogeneity and genomic complexity comparable to many solid tumors^1,2^. Despite major therapeutic advances, including proteasome inhibitors, IMiDs, monoclonal antibodies, and more recently, immunotherapies such as CART cell therapies and T cell engagers, most patients ultimately develop multi-drug refractory disease driven in part by clonal evolution of myeloma cells under treatment pressure^3,4^. Identifying genomic events repeatedly selected by therapy is therefore essential to understand resistance mechanisms driven by tumour biology and to prioritize clinical screening and therapeutic targeting.

We initially applied clonal evolutionary analysis to a large whole-genome sequencing (WGS) rrMM cohort (N=386) and compared clonally-inferred ordering of genomic events with an unrelated newly diagnosed MM (ndMM) WGS cohort (N=198)^1^. This analysis revealed 1p31.1-p12 (hereafter 1pCEN, a region proximal to the centromere) loss of heterozygosity (LOH) as the only common genomic aberration showing marked clonal expansion specifically in rrMM. We validated this observation in three independent longitudinal datasets^5–7^ and then ascertained that the rrMM-enriched 1pCEN event is independent from the established ndMM poor-prognosis locus at 1p32^8^, using the MMRF CoMMpass dataset (N=855)^9^.

## Methods

We re-analyzed WGS from rrMM (N=386) and ndMM (N=198) cohorts previously reported by our group^1^, together with three longitudinal datasets for validation: UK WGS (50 patients, 117 samples)^5^, IFM-2009 WGS (67 patients, 134 samples)^6^ and MMRF CoMMpass WES (66 patients, 129 samples)^7,9^. For hazard-ratio analyses across 1p, we used copy-number segments and Interim Analysis 21 (IA21) release survival data from MMRF CoMMpass (855 newly diagnosed cases)^7^.

Reads were aligned to hg38 and processed under GATK best practices. Somatic SNVs/indels were called and annotated, and allele-specific copy number and tumour purity/ploidy were estimated with Battenberg^10^. Enriched copy number aberration (CNA) regions of gain, LOH and homozygous deletion types were identified by aggregating events across rrMM samples and testing enrichment against a random background using a permutation test (n=1,000) followed by FDR correction (FDR<0.05) as described previously^1^. We utilized high-resolution subclonal heterogeneity analysis to dissect the clonal architecture of MM. Subclonal structure and cancer cell fractions (CCF) were inferred with Battenberg^10^ and multi-dimensional DPClust^11^ as gold-standard methods to determine subclonal structure and evolution^12^. These methods enabled precise estimation of CCF and the identification of subclonal populations^11^. The timing of events across reconstructed phylogenies was obtained via a Plackett-Luce partial-ranking model^13^ in CCF space, enabling chronological ordering (i.e. clonal timing ranking) and assigning timing ranks to recurrent events^14,15^. Fisher’s exact test compared categorical variables; Kaplan-Meier curves, log-rank tests and Cox proportional hazards models were used for survival analysis. All statistical analyses were performed in R (v3.4.3).

## Results and discussion

Chronological ranking of recurrent genetic events (Figure 1A) across rrMM phylogenies (N=386) revealed strong selection and thus clonal expansion of the most recent common ancestor (MRCA) harboring 1pCEN LOH. Its timing-rank dropped nearly four-fold in rrMM when compared with ndMM rankings (timing-rank fold-change = 3.7; P<2.2×10 □^1^ □) (Figure 1B). The 1pCEN locus is adjacent to but is distinct from the prognostic 1p32 LOH and its biallelic events described previously in ndMM. While 1qGain is an early secondary event in ndMM that is further enriched at relapse in frequency^1^, 1pCEN LOH was found to be an enriched CNA (Figure 1C) clearly undergoing MRCA selection in rrMM, indicating a therapy-driven process rather than a simple increase in prevalence. Notably, the intra-tumoral co-occurrence of 1pCEN LOH and 1qGain (hereafter 1pCEN+1q) was significantly enriched in rrMM (co-occurrence OR = 2.3, 95% CI 1.5–3.8, P=2×10□ □) (Figure 1D).

**Figure 1.**
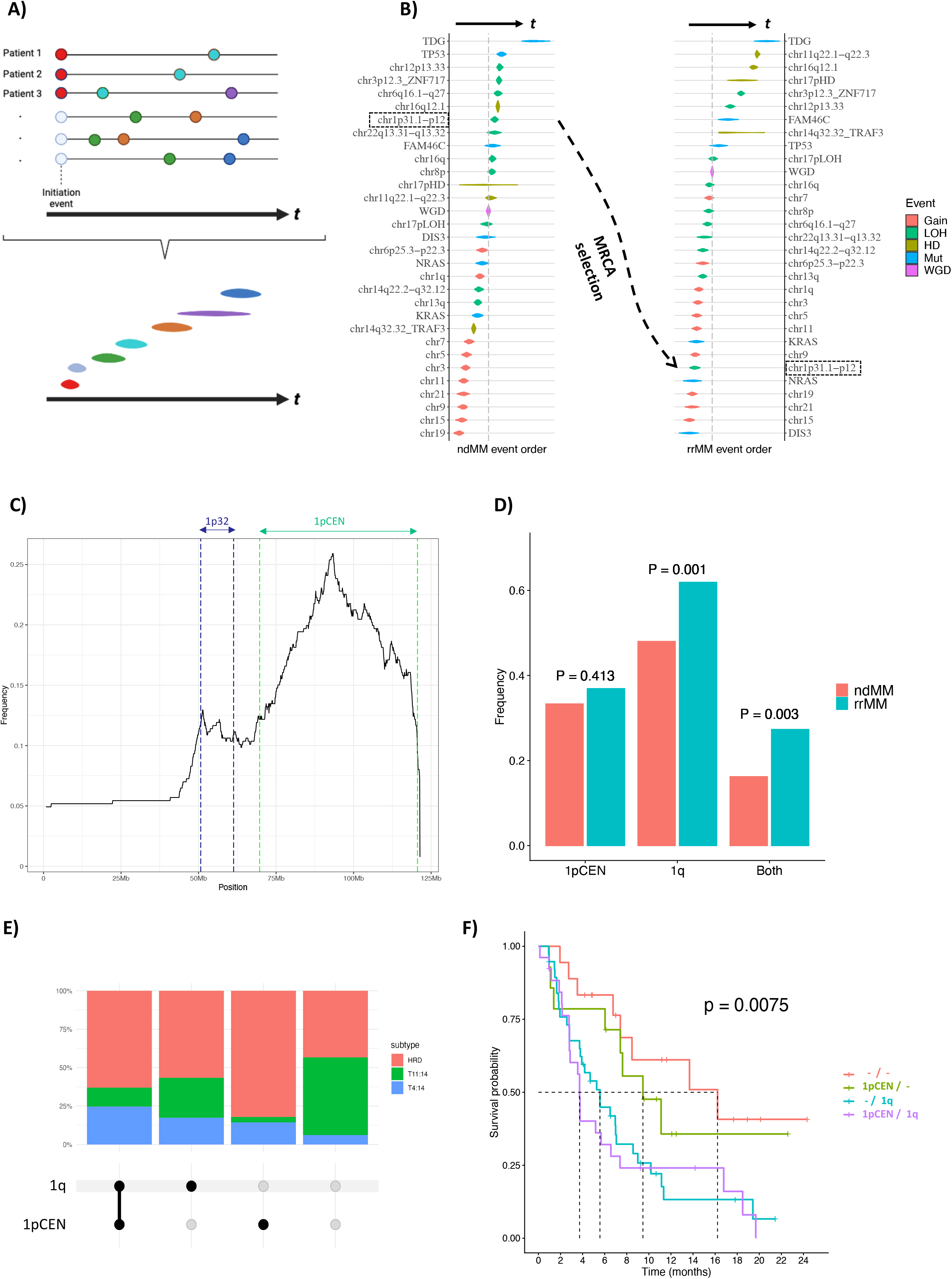
Clonal selection of 1pCEN LOH and enrichment of the poor outcome-associated 1pCEN+1q double-hit in rrMM (N=386). A) Graphical model of the approach used to obtain the most likely order of events based on a partial ranking model of somatic events. Each horizontal violin plot represents the 95% confidence interval of the relative clonal ranking of each event across all tumor phylogenies.Colors are arbitrary and used solely to aid visualisation. B) The identification of 1p31.1-p12 (1pCEN) LOH (dotted black box) as a selected event in rrMM based on tumor phylogenies. Given that the ndMM and rrMM cohorts were unpaired, the chronological order was inferred separately for each cohort. Order of events in the ndMM likely reflect the order of acquisition of somatic events at the start of tumorigenesis while in rrMM the early events are those that were present in the most recent common ancestor (MRCA). Presence of a late ndMM event as an early event in the rrMM cohort is an indication of MRCA selection where the MRCA cell has gained a selective advantage by harboring that event. The 1pCEN LOH event (dashed-border red box) shows a strong MRCA selection signal with nearly a four-fold drop in ranking position and is likely to be associated with relapse/refractory status. C) Frequency pile-up plot for 1p deletion in this dataset delineating the 1p32 region from the enriched 1p31.1-p12 (1pCEN) region in rrMM. D) The frequency distribution of all 1pCEN LOH, all 1qGain and both events in ndMM and rrMM single-timepoint cohorts. E) Upset plot of chromosome 1 copy number event combinations with subtype proportions based on primary events in the rrMM dataset showing the HRD subtype as a suitable subset for survival analysis across the four chromosome 1 categories. F) Kaplan-Meier curves showing the impact of the 1pCEN and 1q single-hits, and 1pCEN+1q double-hit on progression-free survival in the HRD subtype of the largest constituent clinical trial (MM010) in the rrMM dataset.

To assess clinical impact, we examined progression-free survival (PFS) in hyperdiploid tumors from the largest cohort in the rrMM dataset (MM010)^1^ to minimize confounding by primary events (Figure 1E-F). 1pCEN+1q patients had significantly shorter PFS than cases without chromosome 1 aberrations (P=0.0036). This is consistent with previous studies linking either CNA to inferior outcome^16,17^ and reporting poor outcomes in cases with co-occurring 1pCEN and 1q^18–20^, of which one shows the co-occurrence confers even greater risk than either event alone^19^, as recognized in the recent updated IMWG guidance. Because 1p32 deletion is a known poor-prognosis locus in ndMM^8,17^, we tested whether it drove this survival effect. In CoMMpass, the highest deletion frequency across 1p was at 1p22, but the maximal hazard ratio for overall survival was centered at 1p32, recapitulating the established ndMM hazard (Figure 2A). Crucially, excluding rrMM cases with 1p32 LOH and retaining only those with LOH in the 1pCEN region strengthened the adverse PFS signal (P=9.4×10 □□; Supplementary Figure 1), indicating that 1pCEN is an independent determinant of poor outcome in rrMM.

**Figure 2.**
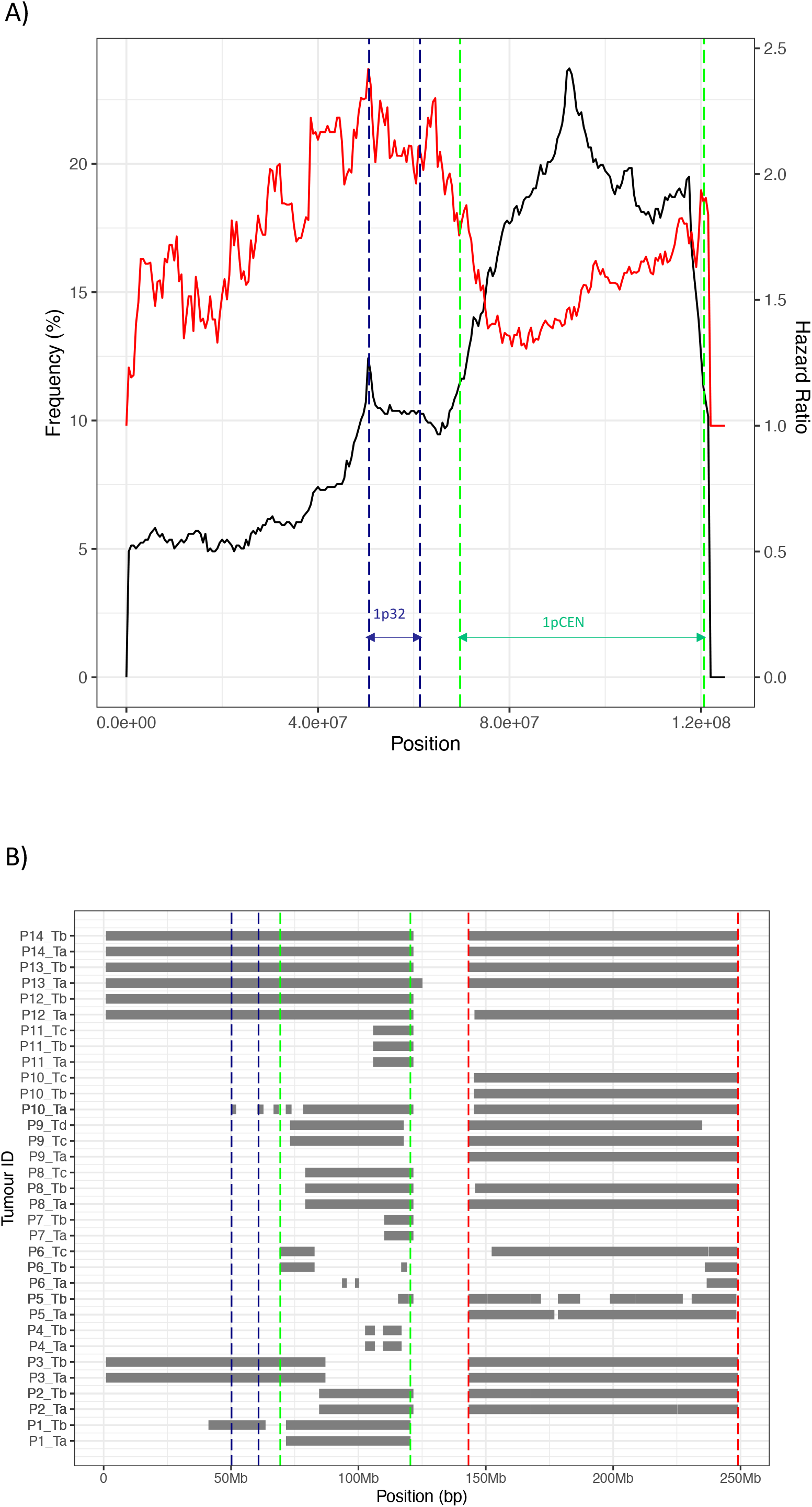
Chromosome 1 deletion events at 1p32 and 1p31.1-p12 (1pCEN) are independent events and associated with different stages of myeloma. A) Deletion and hazard ratio patterns across chromosome 1p in the MMRF CoMMpass ndMM dataset (N=855) showing 1p32 as the main hazard at the newly diagnosed stage. Deletion frequency (black line, left y-axis) and overall survival hazard ratio (red line, right y-axis) for each Mb window in chromosome 1p arm (x-axis). Dashed navy lines show the chr1p32 region, and dashed green lines show the chr1p31 to p12 region (i.e. the RRMM-enriched region 1pCEN). B) Segment pile-up plot displaying co-evolution of LOH across chr1p with 1qGain in the Oxford sequential dataset in patients with the starting sample at the newly diagnosed stage. Navy and green intervals represent LOH at 1p32 and 1p31.3-1p12 (1pCEN) regions. The red interval represents the enriched 1q Gain region. Ta, Tb and Tc represent timepoints 1, 2 and 3 in each patient respectively, with timepoint 1 representing the newly-diagnosed stage in all 14 patients (P1 to P14).

The dissociation between deletion frequency, prognostic hazard, and therapy-enriched 1pCEN highlights distinct biological roles for different 1p sub-regions, mediated by different driver genes and CNAs. In longitudinal analysis of sequential samples from patients with the starting sample at the newly diagnosed stage (three datasets, N=60), almost all patients acquired 1pCEN LOH, either alone or in combination with 1p32 (Figure 2B, Supplementary Figure 2A-B), while isolated 1p32 LOH was observed in only one patient (Supplementary Figure 2B). This evidence supports the role of 1pCEN LOH as the main therapy-selected partner of 1qGain in rrMM. Interestingly, the patients in the 3 datasets received different initial treatment pointing to a therapy-driven but therapy-agnostic drivers residing in the 1pCEN region of the myeloma genome.

We further validated clonal expansion using three longitudinal datasets processed through a unified pipeline. In the UK WGS cohort (50 patients), 1pCEN LOH was present in 20 patients and expanded clonally in 8/20 (40%) with progression (Figure 3A). 1qGain was detected in 36 patients; 22/36 (61%) were clonal at first sampling and 14 showed later subclonal expansion (Figure 3B). The 1pCEN+1q was present in 12/50 (24%): seven cases were clonal for both events at baseline, while five exhibited sequential expansion (three 1pCEN → 1q, two 1q → 1pCEN) (Figure 3B–C). The IFM-2009 and CoMMpass cohorts showed comparable frequencies of 1pCEN+1q (~20–22%). Across all sequential datasets, 1pCEN and 1q displayed consistent clonal expansion with both evolutionary trajectories represented (Figure 3D). A significant proportion of myeloma patients analyzed by us exhibited 1pCEN+1q clonally at diagnosis, suggesting that these patients have high-risk disease pre-treatment. This is consistent with a previous study showing that patients with clonal 1qGain at diagnosis & those with 1qGain clonal expansion at relapse have overlapping outcome profiles^21^, suggesting the equally-aggressive nature of this CNA pre- and post-treatment.

**Figure 3.**
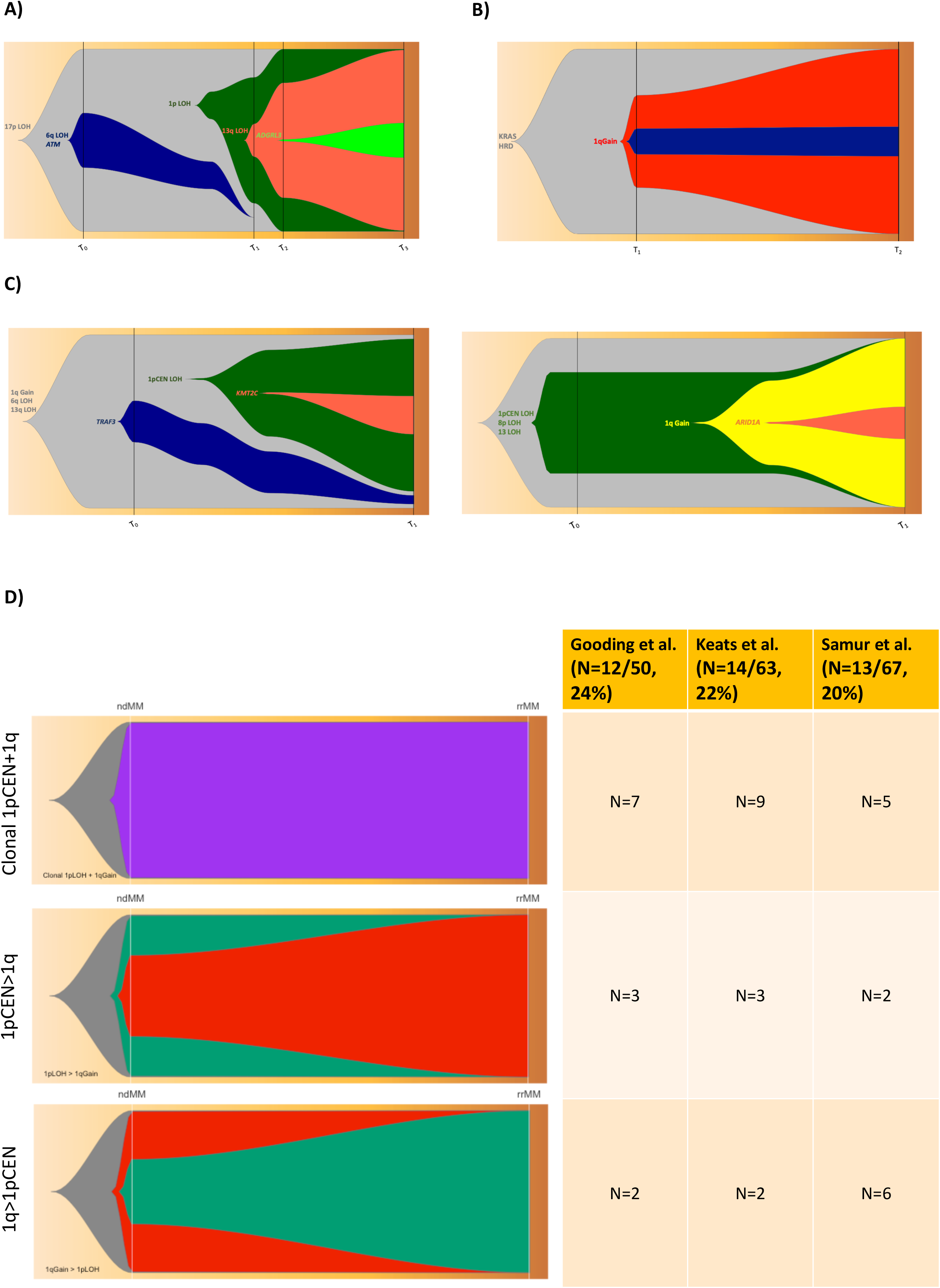
Clonal evolutionary patterns of chromosome 1 copy number aberrations in longitudinal rrMM datasets. A) Fishplot diagram demonstrating clonal expansion of 1p33-p21.1 LOH spanning both 1p32 and 1pCEN in an example patient with four sampling timepoints. The four timepoints represent diagnosis, pre-LEN relapse x2 and post-LEN relapse. B) Fishplot diagram demonstrating clonal expansion of 1q Gain in an example patient with two sampling timepoints. Timepoints represent relapse samples. No driver was identified to map to the navy subclone. C) Fishplot diagram showing expansion of 1pCEN+1q double-hit subclones over sequential samples in both 1q→1pCEN (1p13.2-p12, left) and 1pCEN→1q (1p31.1-p21.3, right) order, in relation to other subclones. D) Fishplot and frequency table diagram for the three evolutionary modes of 1pCEN+1q double-<H1hit (both clonal at T_0_, 1pCEN→1q and 1q→1pCEN) in the three independent patient cohorts with longitudinal sampling.

The recent updated IMWG high risk criteria^8^ specify only the 1p32 region as high risk, as it was developed from ndMM datasets. Integrating our findings across four rrMM datasets and three longitudinal cohorts now also identifies 1pCEN LOH as a recurrent, therapy-enriched lesion that frequently co-occurs with 1qGain to form a double-hit associated with therapy resistance. Crucially, 1pCEN is distinct from the ndMM 1p32 poor-prognosis locus: 1p32 reflects intrinsic aggressiveness at diagnosis, while 1pCEN undergoes selective expansion under therapy and predominates as the partner of 1qGain in rrMM. The observation that both 1pCEN→1q and 1q→1pCEN trajectories occur indicates functional complementarity rather than a fixed order of acquisition. Mechanistically, therapy-associated selection of 1pCEN may reflect loss of tumour suppressors or deletion of regulatory elements altering gene expression that synergize with 1q oncogene dosage (e.g. CKS1B) to drive clonal dominance^22^. More work is needed to dissect the underlying drivers behind this phenomenon. Future research should prioritize fine-mapping of 1pCEN, transcriptomic analysis of double-hit myelomas (e.g. MM.1S cell-line; data not shown), and functional validation of candidate drivers using CRISPR to generate isogenic deletions.

Clinically, screening for 1pCEN and 1qGain at diagnosis or early relapse identifies patients at high risk of evolving the double-hit on treatment. These patients may benefit from ongoing genetic monitoring at subsequent relapses to guide treatment choice. Cost-effective CNA monitoring with for example targeted panels covering 1p32, 1pCEN and 1q^23–25^ could enable serial detection of emerging double-hit clones, as we note that most myeloma FISH probes only target 1p32/1q21 and therefore miss CNAs in this region. Indeed, by using the targeted MGP panel that contains the probes for detecting 1pCEN LOH, out of 21 consecutive rrMM samples from our real-world cohort, we found 1pCEN+1q in 3 cases, demonstrating the routine feasibility of identifying these markers in the clinical samples.

Our study highlights the dynamic nature of MM resistance and provides compelling evidence that the co-occurrence of 1pCEN LOH and 1qGain is a frequent double-hit in the clonal evolution of rrMM under therapeutic selection, but not at the therapy-naive ndMM stage. Screening for this double-hit both at diagnosis and at relapse, particularly in patients whose disease may be acquiring high-risk behavior at recurrent relapses, may aid clinical decision-making^7^. How applicable is our finding in the evolving therapeutic landscape of rrMM driven by immunotherapies? We note that samples in this study were almost all collected in the pre-CAR-T/TCE era. Analyses of genomes of patients treated with CART or TCEs could determine the role played by the 1pCEN+1q double-hit event and more importantly, whether these therapies can suppress the evolution of tumor clones bearing them.

## Acknowledgements

We acknowledge the support of Oxford Translational Myeloma Centre (OTMC) colleagues and funding from OTMC/Oxford University (AT). SG is funded by a Cancer Research UK Fellowship grant RCCCSF□Nov21\100004 and works in an UKRI MRC□funded unit.

## Contributions

The project was conceived and designed by NA-P, SG, and AT. Oversight and management of resources, including data generation, collection, transfer, infrastructure, and data processing, were provided by NA-P, SG, MHK, EFz, SC, AL, EFl, UO, KR, KY, AH, JC, HA-L, NM, MS and AT. Computational and statistical analyses were led by NA-P. Analyses and interpretation were performed by NA-P, SG, MHK, SAH, MS, and AT. Data visualization was performed and structured by NA-P and AT. Supervision and scientific direction were provided by AT. The manuscript was written by NA-P, SG and AT, with contributions and editorial input from MS, AL, EFl, HA-L, and NM. All authors critically reviewed the manuscript and figures.

