## Supplementary Figure 1 for "Sequential acquisition of 1p31.1-p12 LOH and 1q Gain is a common double-hit event in relapsed/refractory myeloma"

**Supplementary Figure 1.** KM plot demonstrating a more pronounced reduction in PFS for 1p31.1-p12 (1pCEN) LOH and 1qGain double-hit (1pCEN+1q) patients after removing those with concurrent 1p32 LOH event.

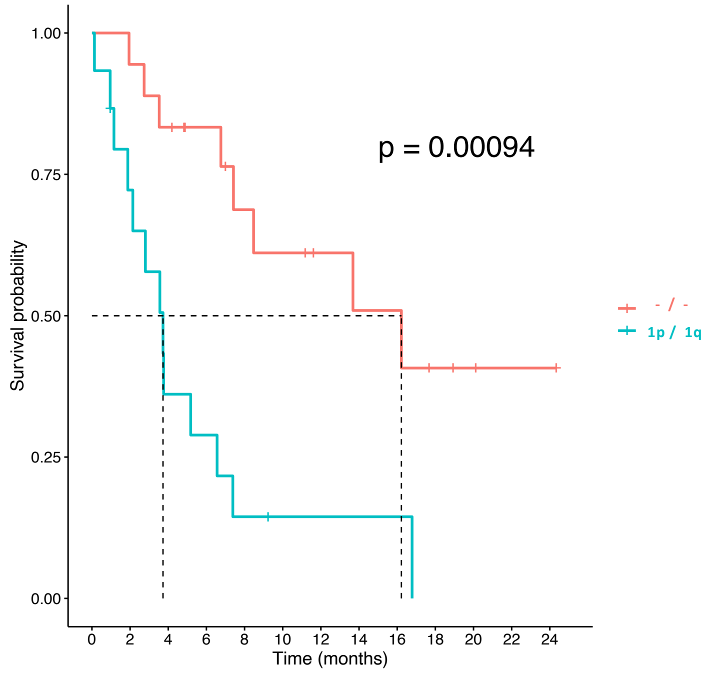
