## Supplementary Figure 2 for "Sequential acquisition of 1p31.1-p12 LOH and 1q Gain is a common double-hit event in relapsed/refractory myeloma"

**Supplementary Figure 2. Co-evolution of 1p LOH with 1q Gain in two additional sequential datasets starting from the newly diagnosed stage.** Segment and frequency pile-up plots for A) DFCI WGS and B) MMRF WES showing co-evolution of 1pLOH and 1q Gain in patients with a sample at diagnosis. Navy and green intervals in 1p represent LOH in 1p32 and the RRMM-enriched region 1p31.3-1p12 (1pCEN). The red interval represents the enriched 1q Gain region. Ta and Tb represent timepoints 1 and 2 in each patient with timepoint 1 representing the newly-diagnosed stage.

A)

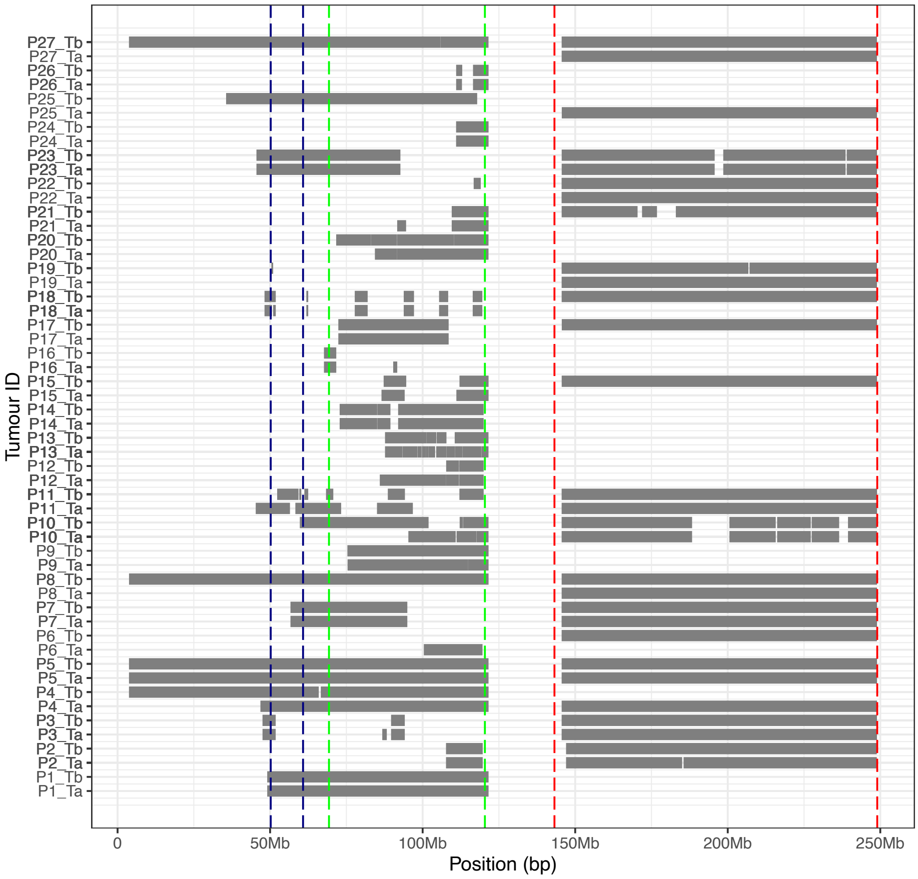


B)

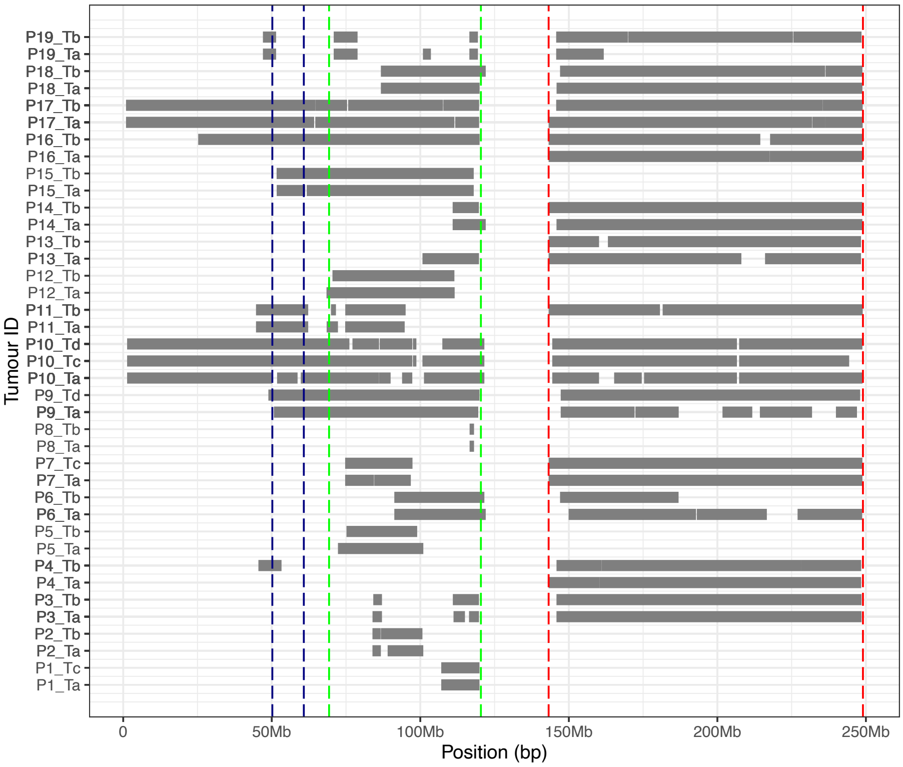
